# Morphometrics reveals complex and heritable apple leaf shapes

**DOI:** 10.1101/139303

**Authors:** Zoë Migicovsky, Mao Li, Daniel H. Chitwood, Sean Myles

**Affiliations:** Department of Plant, Food and Environmental Sciences, Faculty of Agriculture, Dalhousie University, Truro, Nova, Scotia, Canada; Donald Danforth Plant Science Center, St. Louis, MO, USA; Independent researcher, Santa Rosa, CA, USA

**Author notes:** Corresponding author: Zoë Migicovsky.

## Abstract

Apple (*Malus spp*.) is a widely grown and valuable fruit crop. Leaf shape and size are important for flowering in apple and may also be early indicators for other agriculturally valuable traits. We examined 9,000 leaves from 869 unique apple accessions using linear measurements and comprehensive morphometric techniques. We identified allometric variation in the length-to-width aspect ratio between accessions and species of apple. The allometric variation was due to variation in the width of the leaf blade, not length. Aspect ratio was highly correlated with the primary axis of morphometric variation (PC1) quantified using elliptical Fourier descriptors (EFDs) and persistent homology (PH). While the primary source of variation was aspect ratio, subsequent PCs corresponded to complex shape variation not captured by linear measurements. After linking the morphometric information with over 122,000 genome-wide SNPs, we found high narrow-sense heritability values even at later PCs, indicating that comprehensive morphometrics can capture complex, heritable phenotypes. Thus, techniques such as EFDs and PH are capturing heritable biological variation that would be missed using linear measurements alone, and which could potentially be used to select for a hidden phenotype only detectable using comprehensive morphometrics.

## Introduction

Apples (*Malus spp*.) are one of the world’s most widely grown fruit crops, with the third highest global production quantity of over 84 million tonnes in 2014 (1). The shape and size of apple leaves plays an essential role in the growth and development of the tree, and ultimately impact characteristics of the fruit. Apple leaves are generally simple, with an elliptical-to-ovate shape. Previous studies in apple used linear measurements, such as length and width, to quantify leaf shape (2, 3). The length-to-width aspect ratio is a major source of variation in leaf shape. Differing aspect ratios lead to a disproportionate increase or decrease in length relative to width, or allometric variation, in leaves (4, 5). While linear measurements such as leaf length and width are useful, they fail to capture the full extent of leaf shape diversity. Failing to measure leaf shape comprehensively also limits our ability to discern the total underlying genetic contributions.

Elliptical Fourier descriptors (EFDs) are a valuable, well-recognized tool for quantifying the outline of a shape. EFD analysis first converts a contour to a chaincode, a lossless data compression method that encodes shape by a chain of numbers, in which each number indicates step-by-step movements to reconstruct the pixels comprising the shape. A Fourier decomposition is subsequently applied to the chain code, quantifying the shape as a harmonic series. EFDs have been used extensively to quantify leaf shape in diverse species, such as grape (6), tomato (7), and *Passiflora* (8). Previous work used EFDs to assess apple fruit shape (9), but this technique has not yet been applied to apple leaves. A newly developed morphometric technique, persistent homology (PH), provides another method for estimating leaf shape. PH, like EFDs, is normalized to differences in size, but it also could be orientation invariant. PH treats the pixels of a contour as a 2D point cloud before applying a neighbor density estimator to each pixel. A series of annulus kernels of increasing radii are used to isolate and smooth the contour densities. The number of connected components is recorded as a function of density for each annulus, resulting in a curve (a reduced version of persistent barcode) that quantifies shape as topology. The topology-based PH approach can also be applied to serrations and root architecture, allowing the similar framework to be used across different plant structures (10, 11).

Comprehensively measuring leaf shape, using approaches such as EFDs and PH, is important, as shape features may be associated with agriculturally important traits. Leaves are present during the lengthy juvenile phase in apple but fruits appear only on mature trees and thus, leaf traits can enable early selection without the need for genetic markers. In apple, it generally takes 5 years for significant fruiting to occur and any ability to discard trees not possessing a trait of interest earlier in development is extremely valuable (12). There are already several cases of unique leaf characteristics providing an early marker for other genetic differences in apple. For example, the gene underlying red fruit flesh color may lead to anthocyanin accumulation in the leaves, causing red foliage (13, 14) while columnar tree architecture may be accompanied by an increase in leaf number, area, weight per unit area and length-to-width ratio (15). Leaf pH has also been proposed as an early indicator of low acid fruit (16).

In addition to serving as early markers for other traits, leaf shape and size may influence the amount of light a tree receives, and light exposure is crucial for flowering in apple. Light penetration results in higher levels of flowering, while leaf injury or defoliation can reduce flowering (17). Thinning apple trees to a particular leaf-to-fruit ratio is a common practice to attain optimal fruit color and size. Contrastingly, trees with fewer fruit may increase vegetative growth and thus leaf area (18). In previous work, several leaf traits such as area and perimeter were correlated with apple fruit size (19). Clearly, there is an important relationship between the leaves and the fruit, and comprehensively quantifying the variation in leaf shape is a crucial component to understanding this relationship in apple.

Leaves are the main photosynthetic organs of apple, but the genetic basis underlying their shape and size remains unknown. In cotton, a single locus controls the major leaf shapes (20), but in most instances leaf shape appears to be controlled by numerous small-effect loci (5, 21). There are limited examples of genomic analyses of leaf shape in apple, however, a previous bi-parental linkage mapping study found two suggestive quantitative trait loci for leaf size (2). Previous work also measured several leaf traits such as area, perimeter and circularity, in 158 apple accessions. The study linked these measurements with 901 single nucleotide polymorphisms (SNPs) but found no significant genotype-phenotype relationships (19). Thus far, efforts have not been made to estimate the genetic heritability of comprehensive morphometric leaf phenotypes, such as those described using EFDs and PH. It therefore remains unclear to what extent these methods are capturing biologically meaningful, heritable variation.

To fully understand the genetic basis of leaf shape, it is essential to include both linear and morphometric estimates of shape. Decreasing sequencing costs and access to a large and diverse germplasm collection allowed us to analyze approximately 9,000 leaves from over 800 unique accessions which we linked to over 122,000 genome-wide SNPs. We present the first comprehensive analysis of leaf shape in apple, revealing that both accessions and species show allometric variation due to differences in the width of the leaf blade. While the primary axis of variation in apple using EFDs and PH is due to this allometric variation, we find high narrow-sense heritability values even at later principal components, indicating that comprehensive estimates of shape capture heritable variation which would be missed by linear estimates alone.

## Results

### Variation in apple leaf shape

We examined 24 phenotypes related to apple leaf shape and size including length, width, surface area, dry weight, leaf mass per area, within-tree variance, and overall shape estimated using PCs derived from EFD (elliptical Fourier descriptor) and PH (persistent homology) data (see Materials and Methods and Figure 1–2). The sample size and distribution of each phenotype, as well as the raw data, are provided (Figure S1; Table S1).

**Figure 1.**
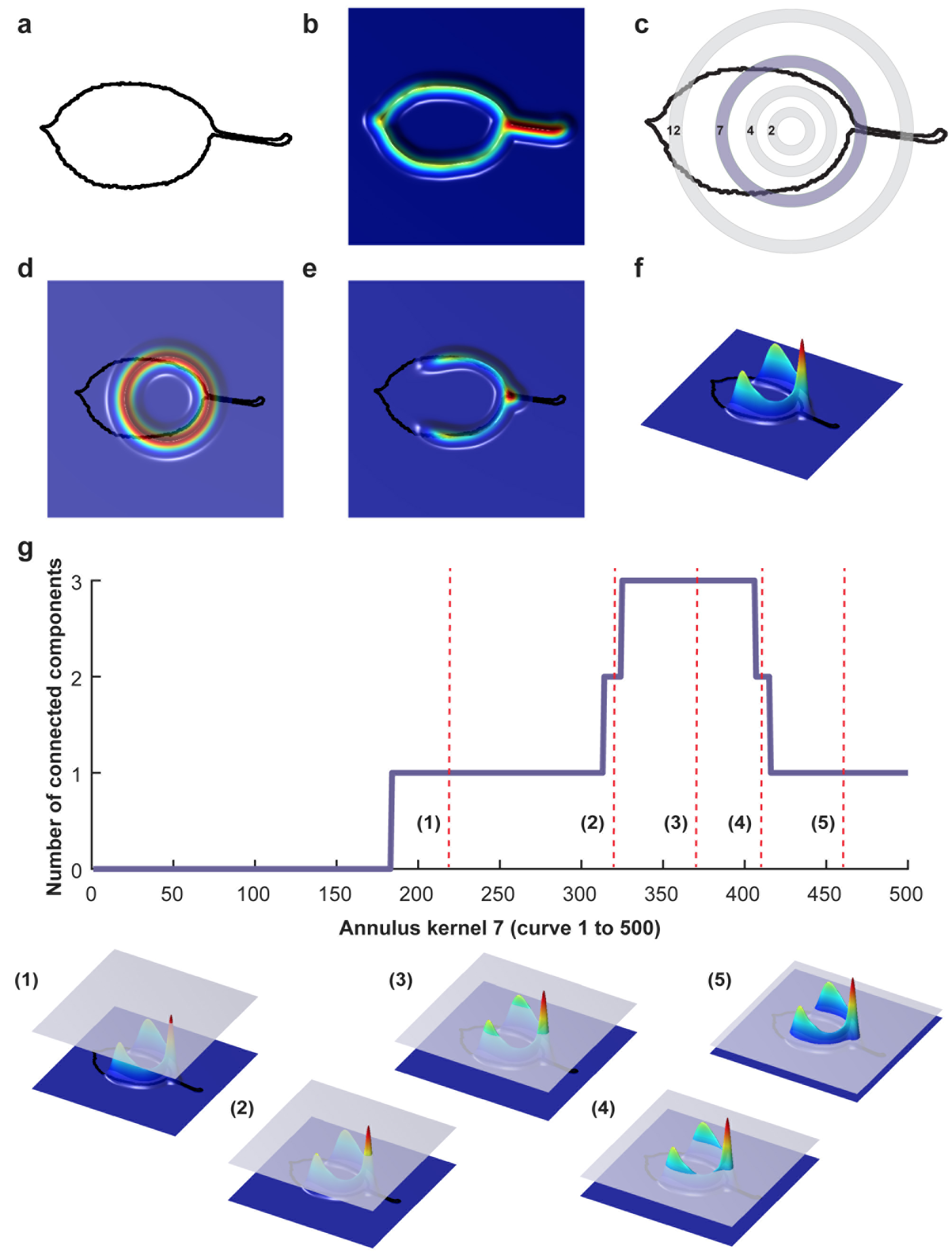
Visualization of persistent homology technique for annulus kernel 7. Binary images were converted into a 2D point cloud (a) which was then normalized using a Gaussian density estimator (b). For each leaf, 16 annulus kernels were used. Annulus kernel 7, indicated in purple (c) is used as an example for this visualization. The density estimator is multiplied by ring 7 (d). The function can also be visualized from the side view (e, f). As a plane moves from top to bottom, the number of connected components is recorded along the curve (g). Below (g) are five visualizations of curves that are represented as red vertical dotted lines in (g).

**Figure 2.**
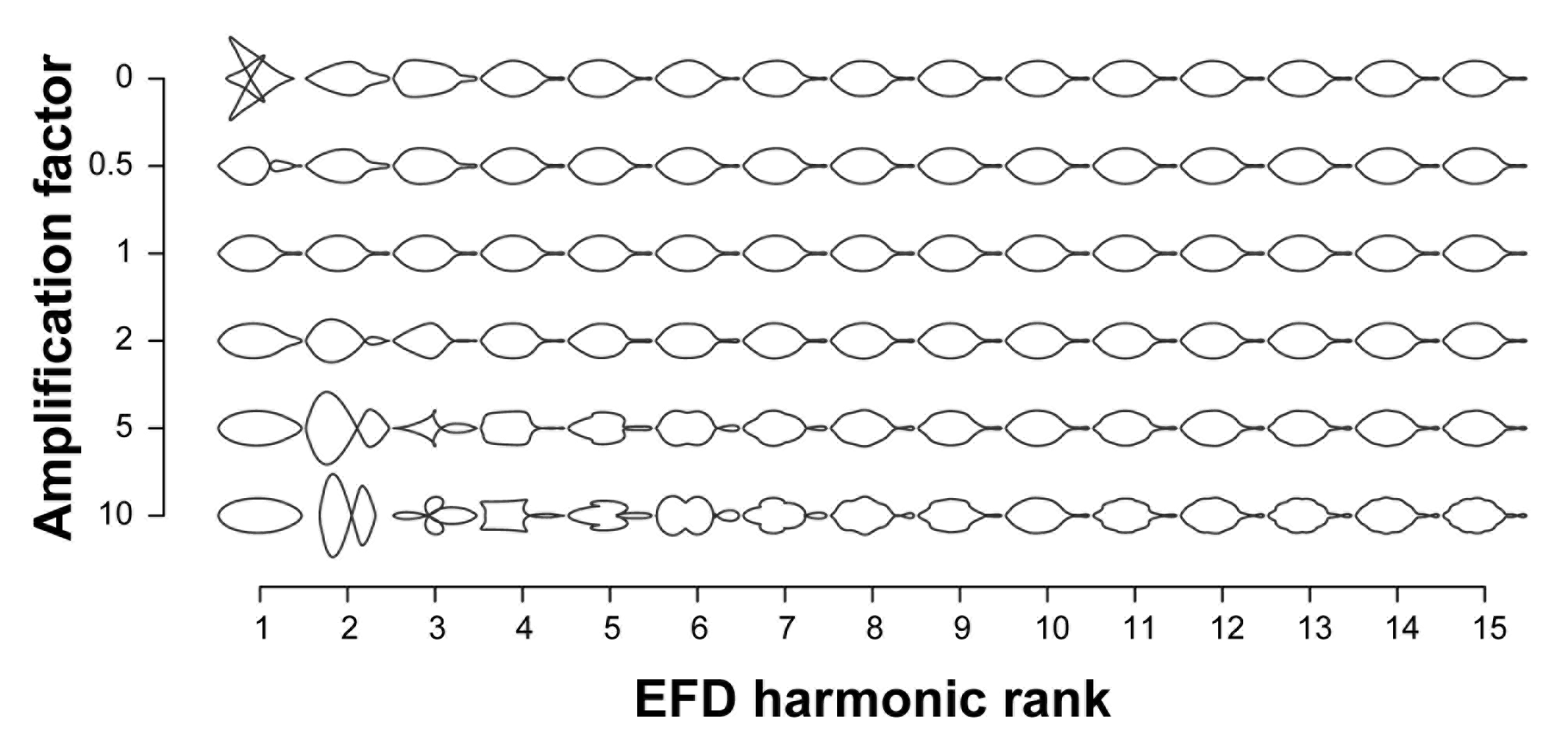
Contribution of elliptical Fourier descriptor harmonics to leaf shape. The leaf shapes depicted are the mean leaf shapes based on all 915 trees. Harmonics 1 to 15 are represented on the x-axis and each harmonic is multiplied by the amplification factor on the y-axis to visualize their contribution to mean leaf shape. An amplification factor of 0 indicates the removal of the harmonic; a factor of 1 results in the normal shape; and values above 1 exaggerate effects to better visualize the harmonic’s contribution to the final shape.

To visualize the primary axes of morphometric variation, we chose a representative leaf from accessions with the minimum and maximum values along the first 5 PCs for EFDs and PH (Figure 3a). The accessions with extreme values along PC1 for both methods are similar. In fact, ‘Binet Rouge’ has the lowest value along PC1 for EFD and PH, with the axis clearly representing a decrease in the length-to-width (aspect) ratio. The annulus kernels most strongly contributing to PH PC1 (Figure S2) provide further evidence that this PC captures variation in aspect ratio. Variation in leaf shape captured by higher-order PCs is more complex and cryptic, and is thus not captured using linear measurements alone. In addition, while the primary axis of variation (PC1) using EFDs and PH may explain similar aspects of leaf morphology, the morphospaces resulting from the two techniques differ (Figure 3b).

**Figure 3.**
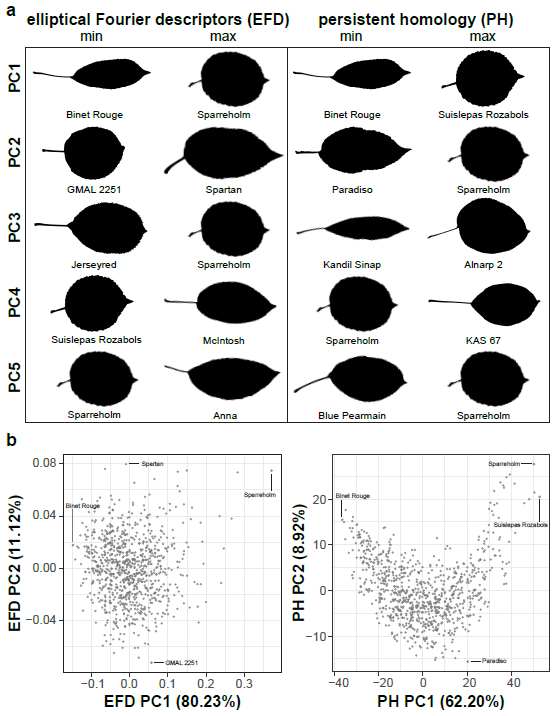
Examples of leaf shape across PCs derived from EFDs and PH. Binary images of leaves from accessions with minimum and maximum values along PCs 1 to 5 for EFD and PH estimates. PCs were calculated using values estimated as the average across 8-10 leaves but only a single representative leaf is displayed. PCs were REML-adjusted based on tree position in the orchard. The accession name is also listed (a). Visualization of PC1 vs PC2 for EFD and PH data. Accession with minimum and maximum values along PC1 and PC2 are indicated (b).

Next, we examined the correlation between all measured traits (Table S3). By assessing the correlation of PCs resulting from a classical morphometric technique such as EFDs with a novel, topology-based morphometric approach like PH, we reveal how complementary the methods are (Figure 4; Figure S3). While there is a highly significant correlation between PC1 for both methods (R^2^ = 0.949, p < 1 × 10^−15^), later PCs are often not significantly correlated, with the most notable exception being EFD PC2 and PH PC3 (R^2^ = 0.432, p < 1 × 10^−15^), although several other PCs also show weak correlations. Thus, while the primary axis of variation (PC1) is consistent and highly correlated between methods, each method captures distinct aspects of leaf morphology in subsequent PCs.

**Figure 4.**
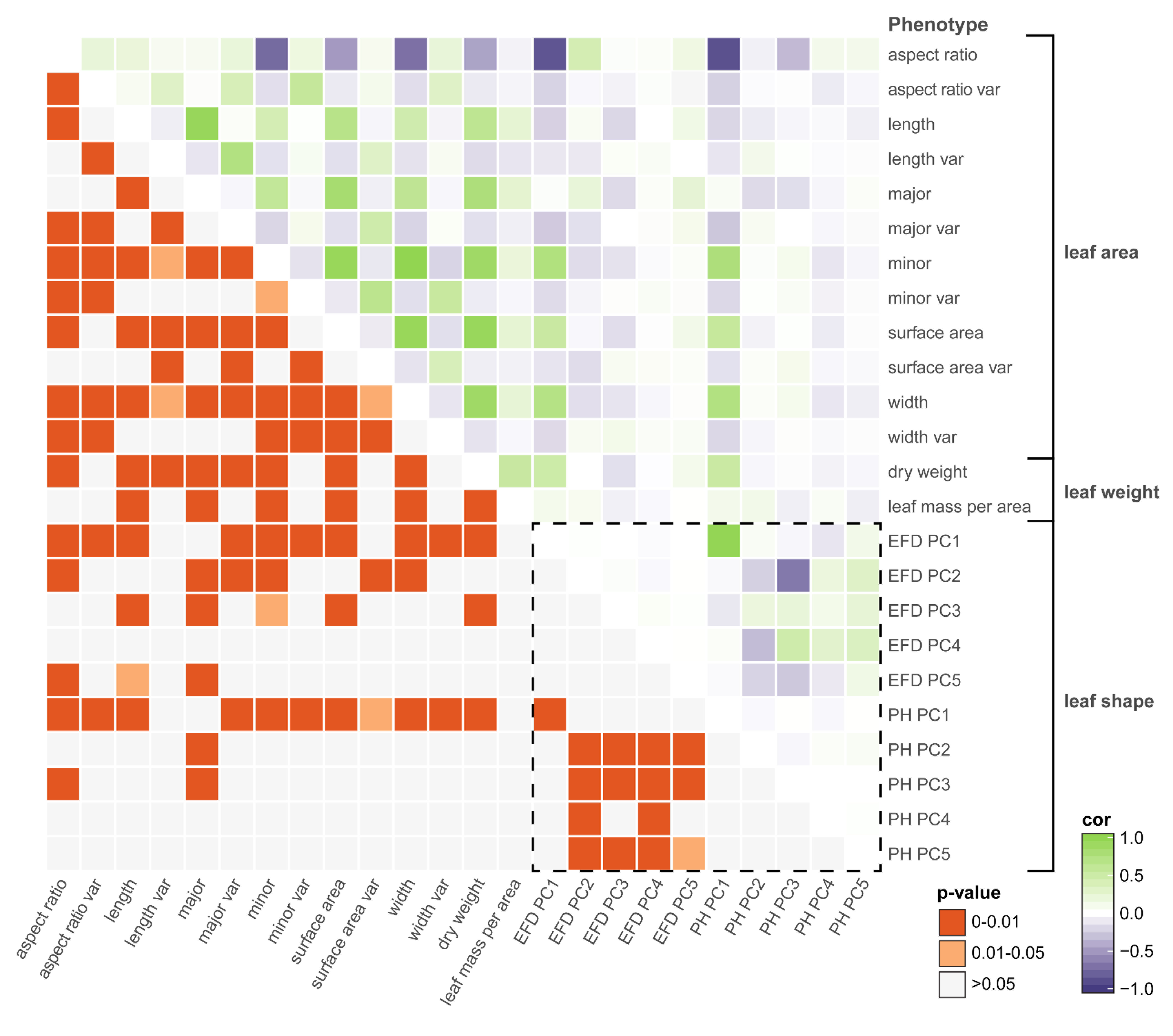
Correlations among leaf phenotypes. Values above the diagonal are colored according to the Pearson’s correlation coefficient, and those below the diagonal indicate Bonferroni-corrected p-values. The box enclosed by the dotted lines include comparisons only between phenotypes captured by comprehensive morphometric analyses.

Many of the leaf phenotypes show a strong correlation with each other (Figure 4). In particular, aspect ratio is highly correlated with PH PC1 (r = −0.878, p < 1 × 10^−15^), EFD PC1 (r = −0.855, p < 1 × 10^−15^) and minor axis (leaf blade width) (r = −0.734, p < 1 × 10^−15^). The correlation between the minor axis of a leaf and surface area (r = 0.939, p < 1 × 10^−15^) is higher than the correlation between the major axis (blade length) and surface area (r = 0.810, p < 1 × 10^−15^). As expected, leaf surface area is also highly correlated with average leaf dry weight (r = 0.934, p < 1 × 10^−15^), indicating that larger leaves are heavier.

### Allometry in apple leaves

The high correlation between aspect ratio and PC1 for both EFD and PH methods indicates that length-to-width ratio is the primary source of variation in apple leaf shape. If there is an allometric relationship between the minor and major axis, and thus, the length and width of a leaf do not increase at equal rates, a slope significantly differing from 1 is expected. We find that the slope between the two measurements is significantly greater than 1 (95% CI = 1.506-1.678, R^2^ = 0.343, p < 1 × 10^−15^), indicating that the minor axis increases at a greater rate than the major axis. While there is no significant correlation between the major axis (blade length) and EFD PC1 (R^2^ = 0.001, p = 1) or PH PC1 (R^2^ = 0.002, p =1), there is a significant correlation for the minor axis (blade width) and EFD PC1 (R^2^ = 0.541, p < 1 × 10^−15^) and PH PC1 (R^2^ = 0.573, p < 1 × 10^−15^) (Figure 5). As PC1 explains 80.23% of the variation in the leaf shape for EFDs, and 62.20% for PH, it is apparent that the width of the leaf blade, and not length, is the major source of leaf shape variation in apple. In fact, the aspect ratio, calculated as the ratio of major axis to minor axis, is even more strongly correlated with EFD and PH PC1, with an R^2^ of 0.732 for EFD PC1 (p < 1 × 10^−15^) and R^2^ of 0.771 for PH PC1 (p < 1 × 10^−15^). Given the significant correlation between EFD PC1 and PH PC1 (Table S3), it is not surprising that aspect ratio is highly correlated with both.

**Figure 5.**
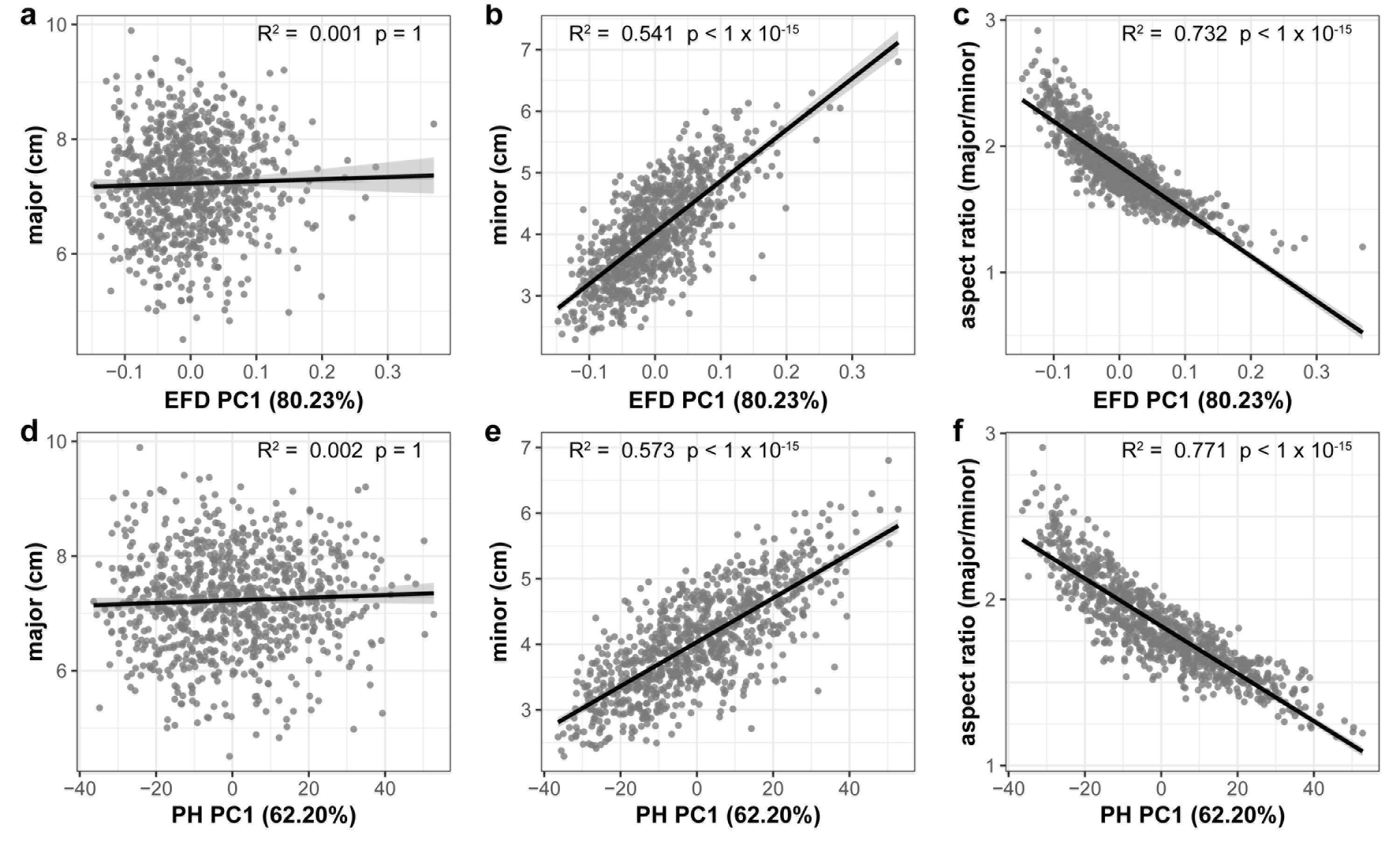
Correlation between the primary axis of variation (PC1) captured using EFD and PH values and leaf shape measures. The EFD PC1 is plotted against the major axis (length of leaf blade) (a), minor axis (width of leaf blade) (b) and aspect ratio (ratio of length-to-width of blade) (c). The PH PC1 is plotted against the same measures in panels d-f. The percent variances explained by PC1, prior to REML-adjustment, is shown in parentheses. All p-values are Bonferonni-corrected based on the number of comparisons in Figure 4. A regression line from a linear model with a shaded 95% confidence interval is also shown.

In addition to variation between accessions, we investigated differences in leaf shape and size between species by comparing *Malus domestica*, the domesticated apple, with its primary progenitor species, *Malus sieversii* (Table S4). PCA of the genome-wide SNP data reveals a primary axis of genetic variation that separates *M. domestica* and *M. sieversii*, although separation is incomplete (Figure 6a). The major axis (p = 0.975) of the leaves does not differ between species (Figure 6b). However, the minor axis (p = 4 × 10^−4^) of *M. domestica* leaves are significantly larger than *M. sieversii* (Figure 6c) and the aspect ratio (p = 0.023) is significantly less (Figure 6d). Thus, there is allometric variation both within (Figure 5) and between (Figure 6) *Malus* species.

**Figure 6.**
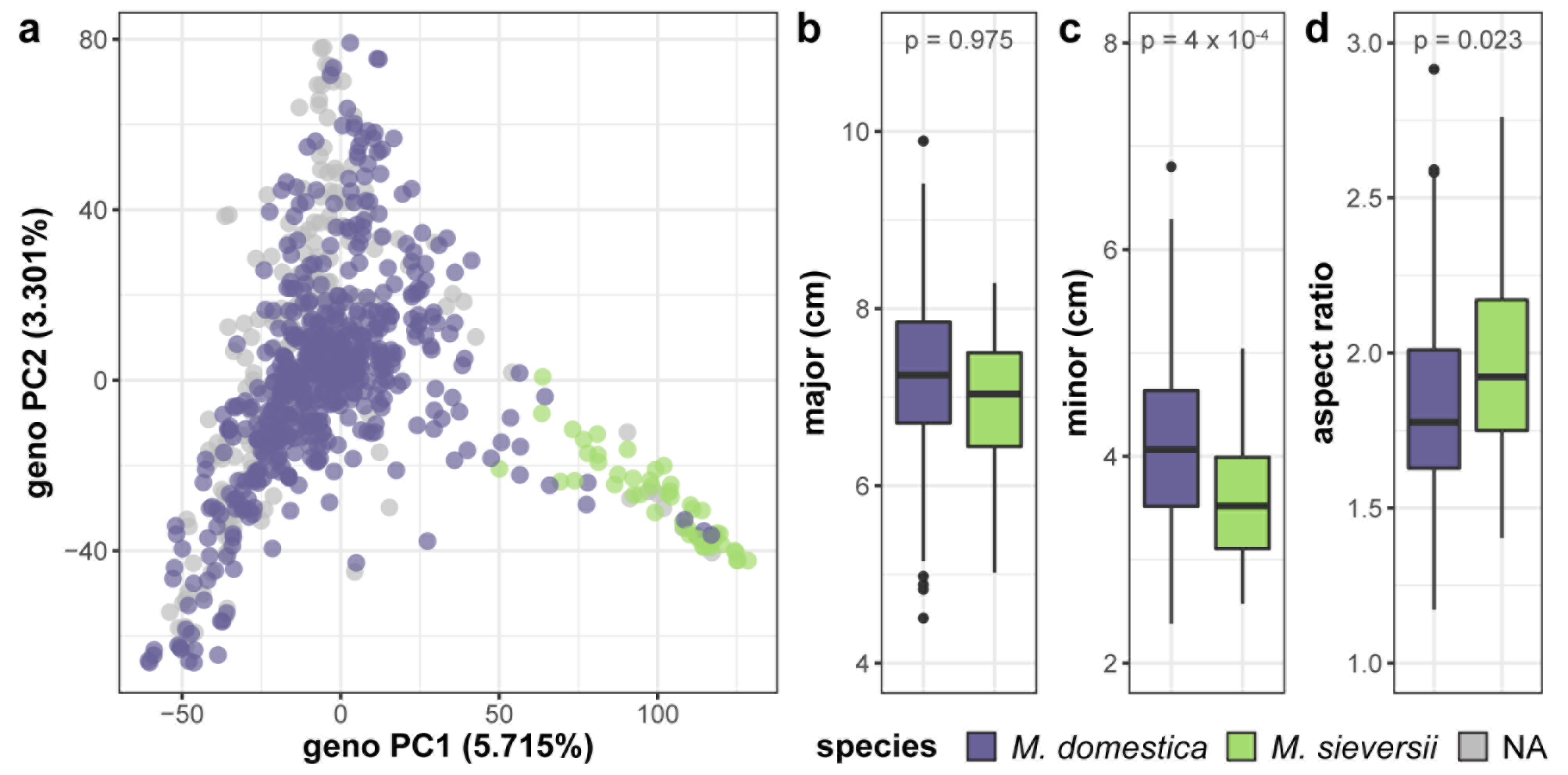
Genetic and phenotypic comparison of the domesticated apple and its wild ancestor. PCs 1 and 2 were derived from 75,973 genome-wide SNPs and samples are labeled as *M. domestica* (purple), *M. sieversii* (green) or unknown (gray). *M. domestica* leaves do not differ from *M. sieversii* leaves along the major axis (b), but they have a larger minor axis (c) and aspect ratio (d). P-values reported are Bonferroni-corrected based on multiple comparisons (Table S4). Species labels are based on USDA classification.

### The genetic basis of leaf shape in apple

GWAS of the 24 leaf phenotypes examined in this study yielded few significant results. We identified 70 significant SNPs representing 5 phenotypes which are reported in Table S5. We examined the regions surrounding significant SNPs for candidate genes using the GBrowse tool (Table S6) (22). We searched within a +/− 5,000 bp window, which should capture any linked causal variation given the rapid LD decay observed in a diverse collection of apples that is largely replicated in the germplasm studied here (23). However, no strong candidate genes were identified.

While GWAS examines the genome for single, large-effect loci, genomic prediction estimates our ability to predict a phenotype using genome-wide marker data. We complimented our GWAS with genomic prediction and observed prediction accuracies (r) ranging from −0.10 to 0.52 (Table S7; Figure S5a). Aspect ratio is the primary source of variation in leaf shape (Fig 5c) and it is also the leaf measurement that had the highest genomic prediction accuracy (0.52). Other phenotypes highly correlated with aspect ratio, such as leaf width (0.51), minor axis (0.49), EFD PC1 (0.48) and PH PC1 (0.47), all had relatively high prediction accuracies. PH PC3 (0.51) was also among the most well-predicted using genetic data.

Similarly, estimates of narrow-sense heritability (h^2^) calculated using GCTA (24) ranged from 0 to 0.75, with the highest heritability observed for aspect ratio (0.75) followed by leaf width (0.71), EFD PC1 (0.71), minor axis (0.69) and PH PC1 (0.65) (Figure 7; Table S8). Heritability estimates were highly correlated with genomic prediction accuracies (Figure S5b, R^2^ = 0.936, p < 1 × 10^−15^), which is not surprising given that both techniques involve predicting a phenotype from genome-wide SNP data. None of the phenotypes measuring variance within the 8-10 leaves sampled had heritability estimates significantly different from 0.

**Figure 7.**
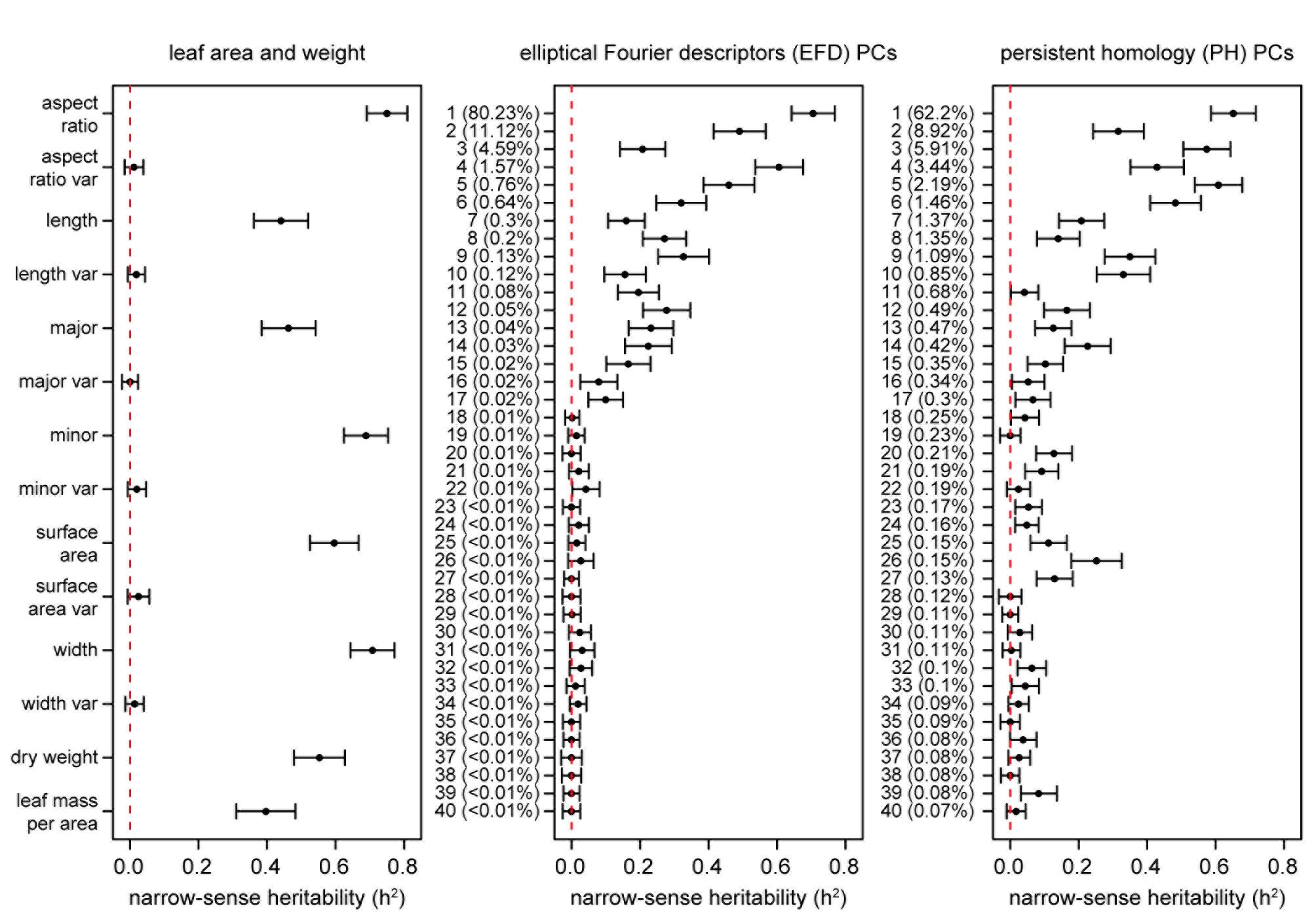
Narrow-sense heritability (h^2^) of leaf phenotypes. Values represent the additive genetic variance (V_g_) divided by the phenotypic variance (V_p_) with a standard error as calculated using GCTA (24). The dotted red lines are found at h^2^ = 0, indicating that none of the phenotypic variance is explained by the genetic data. The proportion of the total phenotypic variance explained by each PC is indicated in parentheses.

While the principal component of variation in leaf shape detected by EFDs and PH is aspect ratio, we were also interested in determining if higher-order PCs, which capture variation not readily visible to the eye, are extracting information that is biologically meaningful. Using genomic prediction and heritability estimates, we found evidence of a genetic basis for these “hidden phenotypes”, which are unmeasurable using linear techniques. For example, the heritability of phenotypes such as PH PC6 (0.48), PH PC9 (0.35), PH PC10 (0.33) and EFD PC9 (0.33) are similar to traditionally measured phenotypes such as leaf length (0.44) and leaf mass per area (0.40). While higher PCs may have relatively high heritability values, after a certain point the values (+/− standard error) overlap with 0, indicating that they are not heritable. The cutoff for morphometric PCs with a heritable genetic basis is approximately PC17. These results suggest that by making use of morphometric techniques that measure shape comprehensively, we are describing biologically meaningful, heritable phenotypes which would be missed by simple measurements such as leaf length, width and surface area.

## Discussion

Leaf shape and size play a crucial role in the growth and development of apple trees, including the fruit. To elucidate the genetic basis of this variation, we quantified leaf shape in apple using traditional linear measurements and comprehensive morphometric techniques. Our work offers the first comparison between the novel topology-based technique, PH, and EFDs, which we find are complementary but distinct methods. For both methods, PC1 was highly correlated with the aspect ratio, thus providing evidence that the primary axis of variation in apple leaf shape can be captured using linear measurements. The minor axis, or width of the leaf blade, was also highly correlated with PC1, while the major axis was not. Thus, variation in the aspect ratio is due to variation in the leaf blade width, not length. Leaf surface area was also more highly correlated with the minor axis than the major axis. Variation in leaf width is therefore essential to both the size and shape of apple leaves, similar to previous work in tomato (25).

The width of the leaf blade is not only the source of variation between apple accessions, but also between *M. domestica* and *M. sieversii*. The presence of the same allometric relationship within and between species suggests that the genetic loci controlling intra-specific leaf shape variation within *M. domestica* may be the same as those controlling the divergence in leaf shape observed between the domesticated apple and its wild ancestor. For example, in birds, while PC1 and PC2 of bill shape explain the majority of variation across 2,000 species, they are also consistently associated with the variation between higher taxa (possessing >20 species) (26). Our results suggest that the increase in leaf size since domestication has not been an overall increase in leaf size but specifically an increase in blade width leading to larger leaves with a reduced length-to-width ratio.

Our work provides evidence that allometry is the primary source of morphometric variation in apple leaves. These findings are consistent with work reported in other species such as tomato, where the length-to-width ratio was the major source of shape variation (>40%) (5). Similarly, work in *Passiflora* and *Vitis* species performed using two independent morphometric techniques identified allometric variation as the primary source of variation in PC1, which explained at least 40% of the variation in leaf shape (8, 27). Thus, linear measurements—in particular aspect ratio—are an important source of information when describing leaf shape. However, linear measurements are not sufficient for capturing the full spectrum of diversity. In our study, PC1 accounts for 62.20% or 80.23% of the variation, depending on the technique used. By simply quantifying apple leaves using linear measurements, we would miss nearly 40% of the variation in some cases. While PC1 is highly correlated with aspect ratio, later PCs represent orthogonal variation that can likely only be captured through morphometric techniques such as EFDs and PH. To fully quantify variation in leaf shape, comprehensive morphometric techniques are therefore essential.

To discern the genetic contributions to leaf shape, we paired both linear and comprehensive morphometric estimates of shape with genome-wide SNP data. There are examples of a simple genetic basis of leaf shape, such as in *Arabidopsis thaliana*, where the *ANGUSTIFOLIA* and *ROTUNDIFOLIA3* independently control leaf width and length (28). In barley, transcript levels of *BFL1* limit leaf width, with overexpression resulting in narrower leaves and loss of *BFL1* function resulting in a reduced length-to-width ratio (29). Using GWAS, we found no robust associations with shape phenotypes, observed a low ratio of significant SNPs to the number of phenotypes examined, and found that significant SNPs were sparsely distributed across multiple chromosomes. In addition, the small number of significant SNPs are likely spurious associations due to poor correction for cryptic relatedness, as evidenced by the QQ plots (Fig S4). These observations suggest that leaf shape is likely polygenic and controlled by a large number of small effect loci, such as in tomato and maize (5, 21). In comparison, GWAS on apple fruit phenotypes, such as color and firmness, have revealed strong associations resulting from a small number of large effect loci (23). However, it is possible that large effect loci were missed in the present study, either because of poor reference genome assembly or inadequate marker density. Improvements in genome assembly and increases in marker number will aid to further reveal the genetic architecture of apple leaf shape variation.

Lastly, we investigated the degree to which leaf shape is heritable and can be predicted using genome-wide SNP data. We find that the genomic prediction accuracies of the primary axes of leaf shape variation are similar to previously reported estimates for fruit width (0.48) and length (0.47), indicating that leaf shape is as heritable as fruit shape (23). In combination with few significant GWAS results, high narrow-sense heritability estimates support a polygenic basis for leaf shape. Aspect ratio was identified as the primary source of variation in leaf shape in apple and had the highest genomic prediction and heritability estimates, indicating that there is a genetic, heritable basis for allometric variation in apple. While we did not detect a genetic basis for leaf shape variation within an accession, we intentionally sampled leaves representing the mean of a tree, and this may have diminished power. Further, although the first 5 PCs for both EFDs and PH explain the majority of the variation in apple leaf shape, most PCs from 1 to 14 have heritability estimates above 0.20 and may still represent crucial differences in leaf shape from an ecological, evolutionary, or agricultural perspective. Thus, while our ability to detect the primary axes of variation in leaf shape using genome-wide data is expected, our observation that higher level PCs are also heritable confirms that these comprehensive morphometric methods capture biologically meaningful variation that would be missed by linear measurements alone.

## Conclusions

It is clear from our work that variation in apple leaf shape and size are under genetic control. Further, high genomic prediction and heritability estimates for higher morphometric PCs indicate that techniques such as EFDs and PH are capturing heritable biological variation that will be missed if researchers restrict leaf shape estimates to linear measurements. Based on these results, it may be possible to perform genomic selection for a phenotype that could only be detected using morphometrics. If a higher order PC was correlated with a trait that was difficult or expensive to measure, assessing leaf shape could potentially be used as proxy for that phenotype, in the same manner that red leaf color can be used to select for red fruit flesh color in apples (13,14). Additionally, a better understanding of the variation in leaf shape and size in apple could ultimately have important implications for canopy management, where light exposure is crucial to flowering (17). Ultimately, through the first in-depth study of leaf shape in apple, we uncover allometry between accessions and species, as well as evidence that complex and heritable phenotypes can be captured using comprehensive morphometric techniques.

## Materials and Methods

### Sample collection

Apple trees in Kentville, Nova Scotia, Canada were budded onto M.9 rootstocks in spring 2012. In the fall, the trees were uprooted and kept in cold storage until spring 2013, when trees were planted in an incomplete block design (see “REstricted Maximum Likelihood (REML)” below). Leaves from over 900 trees were collected from August 24th to September 16th 2015. Ten leaves were collected from each tree. Leaves were flattened and placed to avoid touching, then scanned using Canon CanoScan (LiDE 220) Colour Image Scanners. Leaves were then dried for 48 hours at 65 °C and weighed to estimate the total dry weight (g) for each tree.

### Morphometric analyses

Leaf scans were converted into a separate binary image for each leaf using custom ImageJ macros, which included the ‘make binary’ function (30). A new image file was created for each leaf and named after the tree ID. Images were converted to RGB .bmp files and a chain code analysis was performed using SHAPE (31). The chain code was used to calculate normalized elliptical Fourier descriptors (EFDs) in SHAPE. The normalized EFDs were read into Momocs v1.1.5 (32) in R (33) where harmonics B and C were removed to eliminate asymmetrical variation in leaf shape.

The binary leaf images were also analyzed using persistent homology (PH) (10). To numerically estimate the shape of the leaves using PH, we extracted the leaf contour using a 2D point cloud (Figure 1a). After centering and normalizing the contour to its centroid size, we used a Gaussian density estimator (Figure 1b), which assigns high values (red) to pixels with many neighboring pixels, and low values (blue) to pixels with fewer neighboring pixels. We multiplied the density estimator by an annulus kernel, or ring (Figure 1c), which emphasizes the shape in an annulus at the centroid and is thus invariant to orientation (Figure 1d). The resulting function can also be visualized from the side view (Figure 1e,f). As we moved a plane from top to the bottom, we recorded the number of connected components above the plane, forming a curve. With each new component this value increased, and each time components were merged, it decreased (Fig 1g).

For each leaf, we computed 16 curves corresponding to 16 expanding rings. For computational purposes, each curve is divided into 500 numbers, ultimately resulting in the shape of each leaf being represented by 8,000 (16*500) values.

Only leaves for which both EFDs and PH shape estimations were successfully calculated were included in subsequent analyses. Additionally, only trees with 8-10 leaves were included, as leaves were sometimes removed due to tears, folding, or the absence of a petiole which did not allow for accurate quantification of shape. The final dataset consisted of 915 trees with 8-10 leaves, which included 869 unique accessions and 8,995 leaves.

EFDs and PH values were averaged across leaves from an individual tree. The contribution of EFD harmonics 1 to 15 to the mean leaf shape across all trees was visualized using the ‘hcontrib’ function in the Momocs R package (Figure 2). To allow for discrimination between accessions based on leaf shape, principal component analysis (PCA) was performed using the Momocs ‘PCA’ function (32) for EFDs, and the ‘prcomp’ function in R for PH values, which center but do not scale the data. The resulting PC values were adjusted using REstricted Maximum Likelihood (see below). Subsequently, we identified the accession with the minimum and maximum value along each of the first 5 PCs.

In addition to estimating the contour of the leaf using EFDs and PH, we used several more metrics to describe the leaves. Using ImageJ, we automated the measurement of leaf surface area (cm^2^), length (cm) of the leaf and width (cm) of the leaf as well as major (blade length) and minor (blade width) axes of the best fitting ellipse—which excluded the petiole—through batch processes (30). Throughout the manuscript, we use ‘major’ when referring to the length of the leaf blade, and ‘minor’ when referencing the width of the leaf blade. We also calculated the aspect ratio of the leaf, by dividing the major axis by the minor axis. Additionally, leaf mass per area was calculated for 780 trees where we possessed surface area data for all 10 leaves, by calculating the ratio of dry weight to surface area (g/cm^2^).

While linear phenotypes were calculated as an average value for a particular tree, we also estimated variance within a tree for aspect ratio, length, width, major and minor axis, and surface area. Variance was calculated as the coefficient of variation using the ‘cv’ function in the raster package (34) in R to estimate within-tree variability in leaf size, which is indicated as ‘var’ throughout this manuscript.

### REstricted Maximum Likelihood (REML) adjustment of phenotype data

The orchard sampled in this study is an incomplete block design with 1 of 3 standards per grid. The standards, or “control trees”—‘Honeycrisp’, ‘SweeTango’, and ‘Ambrosia’—are replicated across the grid. Leaves from these trees were sampled multiple times across the orchard, which allowed us to correct for positional effects. Each phenotype was adjusted using a REstricted Maximum Likelihood (REML) model which resulted in one adjusted value per accession, even when multiple trees were measured. The impact of row grid (rGrid), column grid (cGrid) and rGrid × cGrid effects were adjusted for using the following REML model:

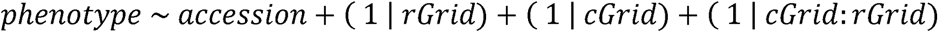

We fit a linear mixed-effects model via REML using the ‘lmer’ function in the lme4 package in R (35) and then calculated the least squares means using the ‘lsmeans’ function in the lsmeans R package (36).

Thus, while the initial phenotype data was collected for 915 trees, following REML adjustment, one value remained per unique accession, resulting in 869 accessions. REML-adjustment was applied directly to all size, weight and variance estimates. For PH and EFDs, we applied the REML following PCA and thus the percent contribution for each PC was calculated using unadjusted values. The adjusted data for all 24 phenotypes are included in Table S1.

### Phenomic analyses

The correlation between leaf phenotypes was calculated using Pearson’s correlation and p-values were Bonferroni-corrected for multiple comparisons. The resulting heatmap was visualized using the ‘geom_tile’ function in ggplot2 in R (37). Next, we examined the leaves for allometry using the ‘SMA’ function in the smartr R package (38) to estimate if the slope between the log-transformed minor and major axis differed from 1.

Accessions were labelled as either *Malus* x. *domestica* Borkh. or *Malus sieversii* Lebed. based on information provided by the United States Department of Agriculture (USDA) Germplasm Resources Information Network website (http://www.ars-grin.gov/) (Table S2). We used a Mann-Whitney U test to test if any phenotypes differed between species and Bonferroni-corrected all p-values for multiple comparisons.

### Genomic analyses

DNA was extracted using commercial extraction kits. Genotyping-by-sequencing (GBS) libraries were prepared using ApeKI and PstI-EcoT221I restriction enzymes according to Elshire, *et al*. (39). GBS libraries were sequenced using Illumina Hi-Seq 2000 technology. Reads which failed Illumina’s “chastity filter” were removed from raw fastq files. Remaining reads were aligned to the *Malus* x. *domestica* v1.0 pseudo haplotype reference sequence (40) using the Burrows-Wheeler aligner tool v0.7.12 (41) and the Tassel version 5 pipeline (42). Tassel parameters included a minKmerL of 30, mnQS of 20, mxKmerNum of 50000000 and batchSize of 20. The kmerlength was set to 82 for ApeKI and 89 for PstI-EcoT22I based on the max barcode size. The minMAF for the DiscoverySNPCallerPluginV2 was set to 0.01. All other default parameters were used. Non-biallelic sites and indels were removed using VCFtools v.0.1.14 (43). VCFs for both enzymes were then merged using a custom perl script, preferentially keeping SNPs called by PstI-EcoT22I at overlapping sites, since those sites tended to be at higher coverage.

Missing data was imputed using LinkImputeR v0.9 (Money et al., Submitted, available: http://www.cultivatingdiversity.org/software.html) with global thresholds of 0.01 for minor allele frequency (MAF) and 0.70 for missingness. We examined depths of 3 to 8 and selected a case for imputation with a max position/sample missingness of 0.70, a minimum depth of 5, and an imputation accuracy of 94.9%. The VCF was converted to a genotype table using PLINK v1.07 (44, 45).

Of the 869 accessions assessed in this study, 816 had genomic data following imputation and filtering and were included in downstream analyses. The resulting genotype table consisted of 816 accessions and 197,565 SNPs. Subsequently, a 0.05 MAF filter was applied using PLINK, after which 128,132 SNPs remained. SNPs with more than 90% heterozygous genotypes were removed. The final genotype table consisted of 816 samples and 122,596 SNPs.

To perform PCA, SNPs were pruned for linkage disequilibrium (LD) using PLINK. We considered a window of 10 SNPs, removing one SNP from a pair if R^2^ > 0.5, then shifting the window by 3 SNPs and repeating (PLINK command: indep-pairwise 10 3 0.5). This resulted in a set of 75,973 SNPs for 816 accessions. PCA was performed on the LD-pruned genome-wide SNPs using ‘prcomp’ in R with data that were centered but not scaled. The first 2 genomic PCs were visualized using ggplot2 in R (37).

We performed a genome-wide association study (GWAS) using the mixed linear model in Tassel (version 5) for each phenotype, adjusting for relatedness among individuals using a kinship matrix as well as the first 3 PCs for population structure (46, 47). The threshold for significance was calculated using simpleM (48, 49) which estimates the number of PCs needed to explain 0.995 of the variance, or the number of independent SNPs. The inferred Meff used to calculate the significance threshold was 91,667 SNPs.

We searched the regions surrounding any significant GWAS SNPs using the Genome Database for Rosaceae GBrowse tool for *Malus* x. *domestica* v1.0 p genome (22). We used a window of +/− 5,000 bp (10 kb) surrounding the significant SNP to check for genes, and when identified, we used the basic local alignment search tool (BLAST) from NCBI to search for the mRNA sequence and reported the result with the max score (50).

Genomic prediction was performed using the ‘x.val’ function in the R package PopVar (51). The rrBLUP model was selected and 5-fold (nFold=5) cross-validation was repeated 3 times (nFold.reps=3) with no further filtering (min.maf=0) from the set of 122,596 SNPs used for GWAS. All other default parameters were used. In addition to performing genomic prediction on the main 24 phenotypes examined in this study, we performed genomic prediction on all 40 PCs for EFDs and on the first 40 PCs for PH values. We also used the ‘rnorm’ function in R to generate 1,000 random phenotypes with a mean of 0 and a standard deviation of 1, and performed genomic prediction using these random phenotypes to obtain the range of genomic prediction accuracies one can expect at random. Lastly, we used genome-wide complex trait analysis (GCTA) v.1.26.0 which estimates the genetic relationships between individuals based on genome-wide SNPs and uses this information to calculate the variance explained by these SNPs. The ratio of additive genetic variation to phenotypic variance is used to calculate narrow-sense heritability (h^2^), or SNP heritability, of a trait (24). We used GCTA to estimate heritability for each phenotype, including the first 40 PCs for EFD and PH. We also estimated the correlation between genomic prediction accuracy (r) and narrow-sense heritability (h^2^) using a Pearson’s correlation.

## Acknowledgments

This article was written, in part, thanks to funding from the Canada Research Chairs program, the National Sciences and Engineering Research Council of Canada and Genome Canada. Z.M. was also supported in part by a Killam Predoctoral Scholarship from Dalhousie University.

We would like to acknowledge Gavin Douglas and Sherry Fillmore for their help setting up the statistical design of the orchard and SNP-calling pipeline. We also thank all past and present members of the Myles Lab for their work in maintaining the apple orchard.

## Supplementary Material

**Figure S1. Distribution of leaf phenotypes following REML-adjustment.** N is equal to the total number of unique samples.

**Figure S2. Visualization of contributions of each ring to PH PC1.** Rings 6, 7 and 16 contribute the most to leaf shape according to PH PC1. The placement of each ring is visualized on a leaf representing the minimum and maximum value along PC1 (a). The contribution to PC1 of each of the 16 rings is also shown (b).

**Figure S3. Comparison of morphometric EFD and PH PCs 1 to 5.** Correlation between first 5 PCs, estimated using Pearson’s correlation, including R^2^ and Bonferroni corrected p-values based on Figure 4/Table S3.

**Figure S4. GWAS results for all 24 leaf phenotypes examined.** Manhattan and QQ plots are included for each phenotype. The QQ-plot shows both the results of a naive GWAS (Pearson correlation) and the results from applying the mixed model. P-values are log-transformed and the threshold for significance is simpleM-corrected and indicated by a horizontal dotted line. Chromosome R indicates SNPs found on contigs unanchored to the reference genome.

**Figure S5. Genomic prediction accuracy (r) (a) and correlation between genomic prediction results and narrow-sense heritability estimates (h^2^) for all leaf phenotypes (b).** Genomic prediction accuracies represent the average correlation (+/− standard deviation) between observed and predicted phenotype scores, based on 5-fold cross-validation with 3 iterations. Dotted red lines indicate the minimum and maximum prediction average accuracy (r) achieved using 1,000 randomly generated phenotypes. The percent variance explained by each PC was calculated prior to REML-adjustment and is indicated in parentheses.

**Table S1. All leaf phenotypes assessed in apple, following REML-adjustment.** Accessions are identified by their unique “apple id”. Further information about these accessions is available in Table S2.

**Table S2. Metadata for all accessions assessed in this study.** In addition to the unique numeric apple_id, we report the Germplasm Origin (where budwood was obtained from) and Species (*Malus domestica*/*Malus sieversii*).

**Table S3. Correlation between leaf phenotypes as well as Bonferroni-adjusted p-values.** Pearson’s product moment correlation coefficients are reported. These results are visualized in Figure 2.

**Table S4. Comparison of leaf phenotypes between accessions based on metadata.** Bonferroni-adjusted p-values resulting from a Mann-Whitney U test estimating the difference between accessions based on species (*Malus domestica*/*Malus sieversii*) for the leaf phenotypes examined.

**Table S5. Positional information for significant GWAS results.** Additional information about significant SNPs are included such as p-value, marker R^2^, minor and major allele, minor and major effect and MAF.

**Table S6. Genes found within +/− 5 kb of SNPs with significant associations to phenotypes from GWAS.** Results are listed according to the Genome Database for Rosaceae GBrowse (accessed January 27 2017). Overlapping mRNA, length, contig, GO category, GO term accession, GO term name, InterPro Term, InterPro Description and NCBI sequence with Max Score when BLASTed using NCBI are reported.

**Table S7. Genomic prediction accuracies (r) for leaf phenotypes.** r_avg represents the average correlation between observed and predicted phenotype scores, based on 5-fold cross-validation with 3 iterations. The standard deviation (r_sd) is also reported.

**Table S8. Narrow-sense heritability (h^2^) for leaf phenotypes.** h^2^ represents the genetic variance (V_g_) divided by the phenotypic variance (V_p_). The standard error (SE) is also reported. These results are visualized in Figure 7.

